# CISD3 is required for Complex I function, mitochondrial integrity, and skeletal muscle maintenance

**DOI:** 10.1101/2023.06.03.543558

**Authors:** Henri-Baptiste Marjault, Ola Karmi, Linda Rowland, Thi Thao Nguyen, DeAna Grant, Camila Manrique-Acevedo, Rachel Nechushtai, Ron Mittler

## Abstract

Mitochondria play a central role in muscle metabolism and function. In skeletal muscles, a unique family of iron-sulfur proteins, termed CISD proteins, support mitochondrial function. The abundance of these proteins declines with aging leading to muscle degeneration. Although the function of the outer mitochondrial proteins CISD1 and CISD2 has been defined, the role of the inner mitochondrial protein CISD3, is currently unknown. Here we show that CISD3 deficiency in mice results in muscle atrophy that shares proteomic features with Duchenne Muscular Dystrophy. We further reveal that CISD3 deficiency impairs the function and structure of skeletal muscle mitochondria, and that CISD3 interacts with, and donates its clusters to, Complex I respiratory chain subunit NDUFV2. These findings reveal that CISD3 is important for supporting the biogenesis and function of Complex I, essential for muscle maintenance and function. Interventions that target CISD3 could therefore impact muscle degeneration syndromes, aging, and related conditions.

## INTRODUCTION

NEET or CDGSH proteins belong to a unique class of highly conserved iron-sulfur [2Fe-2S] proteins^1–3^. Their iron-sulfur binding domain, coordinated by 3Cys-1His (part of the conserved CDGSH domain; <u>C</u>-X-<u>C</u>-X2-(S/T)-X3-P-X-<u>C</u>-D-G-(S/A/T)-<u>H</u>), is redox-sensitive and involved in many different [2Fe-2S] cluster, and/or electron, transfer reactions^4–8^. In humans three different genes encode for NEET/CDGSH proteins (termed <u>C</u>DGSH Iron <u>S</u>ulfur <u>D</u>omain-containing, or CISD proteins). These are: CISD1 or mitoNEET, localized to the outer mitochondrial membrane, CISD2 or NAF-1, localized to the outer endoplasmic reticulum (ER), mitochondria, and ER-mitochondria associated membranes (MAM), and CISD3 or MiNT, localized inside the mitochondria. Although multiple studies have shown that CISD1 and CISD2 are involved in the progression of different human pathologies, including cancer, diabetes, neurodegeneration, obesity, and cardiovascular disease^1–3,9–17^, as well as the genetic disease Wolfram Syndrome 2 (caused by CISD2 deficiency)^18–20^, very little is known about the function of CISD3.

In contrast to CISD1 and CISD2 that are symmetric homodimers attached to a membrane (each monomer containing one [2Fe-2S] cluster), CISD3 is a soluble monomeric protein that folds into a pseudo dimer that contains two, not perfectly symmetric, [2Fe-2S] clusters^1–4,21^. Recent studies support a model in which CISD3 can transfer its [2Fe-2S] clusters from within the mitochondria to CISD1, that is outside the mitochondria, and that CISD1 can transfer these clusters to CISD2 and/or Anamorsin, constituting a [2Fe-2S] shuttle from within the mitochondria to the cytosol^22–25^. In support of this model are the findings that disruption of any one of the three CISD proteins in cancer cells causes an elevation in the levels of mitochondrial labile iron and reactive oxygen species (ROS), that is accompanied by the activation of a ferroptotic and/or apoptotic cell death pathway^2,4, 25–27^. A recent study has also revealed that targeting CISD3 function in cancer cells induces glutaminolysis and causes cystine deprivation-induced ferroptosis^28^. Interestingly, the protein levels of CISD3 (as well as CISD1 and CISD2) are elevated in many different cancer cells, to prevent the activation of ferroptosis and/or apoptosis that may be triggered due to the overaccumulation of ROS and/or labile iron in mitochondria (part of the iron/ROS ‘addiction’ of many cancers) ^2,29,30^.

Studies of CISD3 function in *Caenorhabditis elegans* (*C. elegans*) revealed that this protein is required for maintaining normal germline structure and function by regulating physiological germline apoptosis (through the canonical programmed cell death pathway)^31,32^. CISD3 mutations in *C. elegans* consequently result in germline abnormalities, associated with an irregular stem cell niche, and disrupted formation of bivalent chromosomes^31^. As the phenotypes associated with CISD3 in *C. elegans* were also found to be linked with mitochondrial dysfunction, including disruption of the mitochondrial network within the germline^31,32^, they further support a role for CISD3 in maintaining and/or promoting mitochondrial function in cells with a high energy demand and/or ROS levels (such as stem cells). Remarkably, although CISD3 is conserved in animal cells, some fungi, and bacteria, it does not appear to be present in higher plants^33,34^.

Although the role of CISD3 in cancer cells and *C. elegans* has been previously studied^4,22,28,31^, very little is known about the function of CISD3 in the development and physiological maintenance of animal cells and tissues. To identify a role for CISD3 in animal development and normal physiological functions, we generated a transgenic global deletion (knock-out; KO) mice model for CISD3 (*Cisd3^-/-^*). Our analysis of *Cisd3^-/-^* mice revealed that CISD3 is essential for the maintenance and function of skeletal muscles, and that deficiency in CISD3 results in muscle atrophy that shares proteomic features with Duchenne Muscular Dystrophy (DMD). In addition, we show that deficiency in CISD3 impairs the function and structure of skeletal muscle mitochondria, and that CISD3 interacts with, and donates its clusters to, Complex I mitochondrial respiratory chain subunit NDUFV2 (NADH:Ubiquinone Oxidoreductase Core Subunit V2; encoded by the nuclear gene *NDUFV2*). These findings reveal that CISD3 has an important function within the mitochondria, supporting the biogenesis/function of Complex I, and that this function is essential for muscle maintenance and function. Interventions that target CISD3 function in muscles could therefore impact muscle degeneration syndromes, aging, and related conditions.

## RESULTS

### Characterization of CISD3 knockout mice

C57BL/6J wild type (WT) and *Cisd3^-/-^* male and female mice, were grown under controlled conditions for 44-47 weeks. Compared to WT, *Cisd3^-/-^* male and female mice were significantly smaller (Figs. 1c, 1d, S1) and had a life span of about 60 weeks (about half of WT mice). As the CISD protein family is proposed to be functionally linked^2,22,25^, we tested the expression of all CISD proteins in the quadriceps muscle of WT and *Cisd3^-/-^* mice. This analysis confirmed that *Cisd3^-/-^* mice lacked the CISD3 protein and revealed that the deficiency in CISD3 caused a significant decrease in the levels of CISD1, as well as a significant increase in the levels of CISD2 (Fig. 1e). This finding suggests that the CISD protein network is linked and that changes in the levels of one CISD protein (*i.e.,* CISD3) alters the levels of the two other members of this family (*i.e.,* CISD1 and CISD2).

**Fig. 1:**
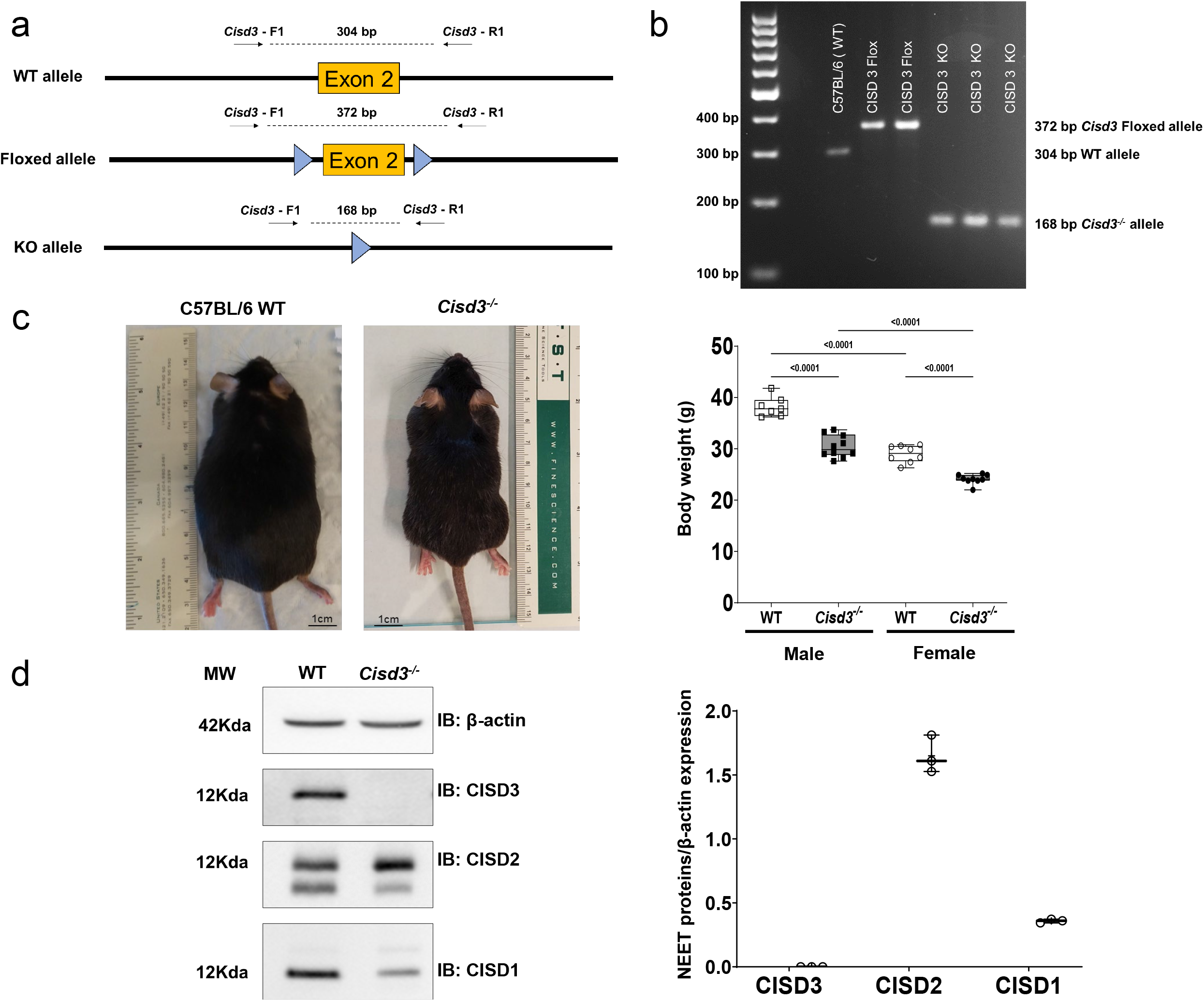
Generation and characterization of *Cisd3^-/-^* mice. **a** Targeting of exon 2 of *Cisd3* using a flox strategy. **b** PCR analysis of wild type (WT), floxed, and the knockout (KO) alleles of *Cisd3* in transgenic mice. **c** Representative images of a WT and a *Cisd3^-/-^* mice (left) and a bar graph showing the weight of WT and *Cisd3^-/-^* mice at age 44-47week-old (right; Fig. S1). **d** Representative protein blots (left) and a bar graph showing quantitative analysis (right) for the levels of CISD1, CISD2, and CISD3 proteins in WT and *Cisd3^-/-^* mice. Results are shown for male and female separately and presented as mean ± SD of 6 different animals (3 different males and 3 different females) from each group (WT and *Cisd3^-/-^* mice). Two-way ANOVA followed by a Tukey test was used to calculate statistical significance. White box and white square, WT male mice; Gray box and black square, *Cisd3^-/-^* male mice; white box and white circle, WT female mice; Gray box and black circle, *Cisd3^-/-^* female mice. Abbreviations: CISD, CDGSH Iron Sulfur Domain; KO, knock out; WT, wild type.

### Deficiency in CISD3 affects skeletal muscle structure and function

When handling the *Cisd3^-/-^* mice, it was found that compared to WT, *Cisd3^-/-^* male and female mice were physically weaker and had a reduced grip strength. We therefore measured their ability to hang from a wheel or a rod^35^. Compared to male and female WT mice, both male and female *Cisd3^-/-^* mice displayed a lower ability to hang from a wheel or a rod (Fig. 2). This finding suggested that compared to WT, the muscle strength of male and female *Cisd3^-/-^*mice is weaker and does not allow them to hang from a wheel or a rod for the same length of time as WT mice can.

**Fig. 2:**
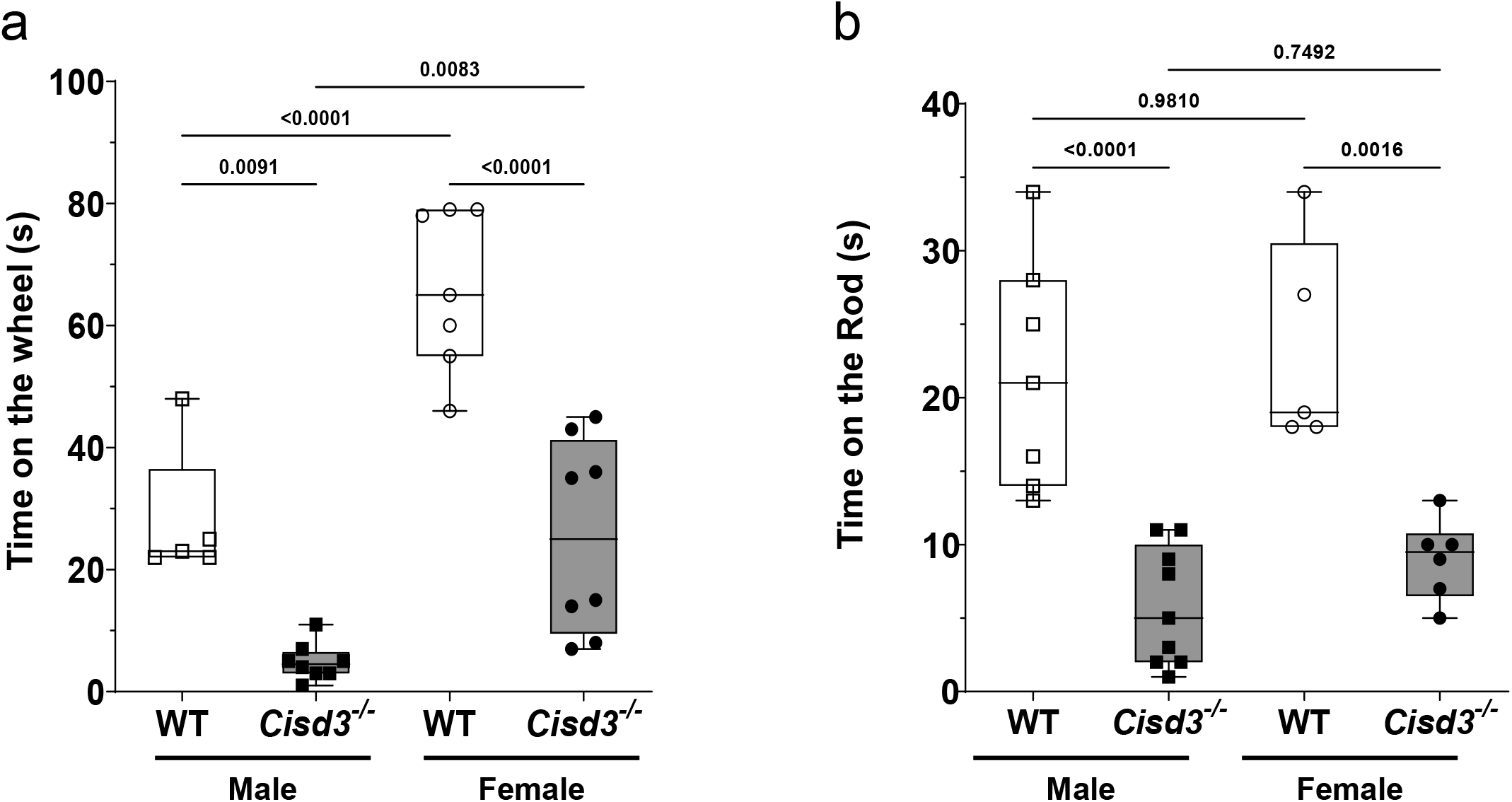
*Cisd3^-/-^*mice display reduced muscle strength. **a** and **b** Bar graphs showing the amount of time wild type (WT) or *Cisd3^-/-^* mice can hang from a wheel (a) or a rod (b). Results are shown for male and female separately and presented as mean ± SD of 6 different animals (3 different males and 3 different females) from each group (WT and *Cisd3^-/-^* mice). Two-way ANOVA followed by a Tukey test was used to calculate statistical significance. White box and white square, WT male mice; Gray box and black square, *Cisd3^-/-^* male mice; white box and white circle, WT female mice; Gray box and black circle, *Cisd3^-/-^* female mice. Abbreviations: CISD, CDGSH Iron Sulfur Domain; KO, knock out; WT, wild type.

To study the effect of CISD3 deficiency on skeletal muscle of mice, we focused on the quadriceps muscle (Fig. 3a) and compared its cellular and subcellular structure between 44-47-week-old WT and *Cisd3^-/-^* male and female mice using light and transmission electron microscopy (TEM). Staining of muscle sections with H&E did not reveal significant differences between WT and *Cisd3^-/-^* mice; except for a larger number of centralized nuclei in muscle cells of male *Cisd3^-/-^* mice compared to male WT mice (Figs. 3b, S2). TEM analyses of muscle tissue from male and female WT and *Cisd3^-/-^* mice revealed that the muscular sarcomere (*i.e.,* the distance between the muscle Z-lines) of male and female *Cisd3^-/-^* mice is shorter compared to that of WT mice (Fig. 3c). A shorter Z-line distance was previously shown to correlate with muscle atrophy^36,37^ supporting our muscle strength rod and wheel hanging test results of *Cisd3^-/-^*mice (Fig. 2).

**Fig. 3:**
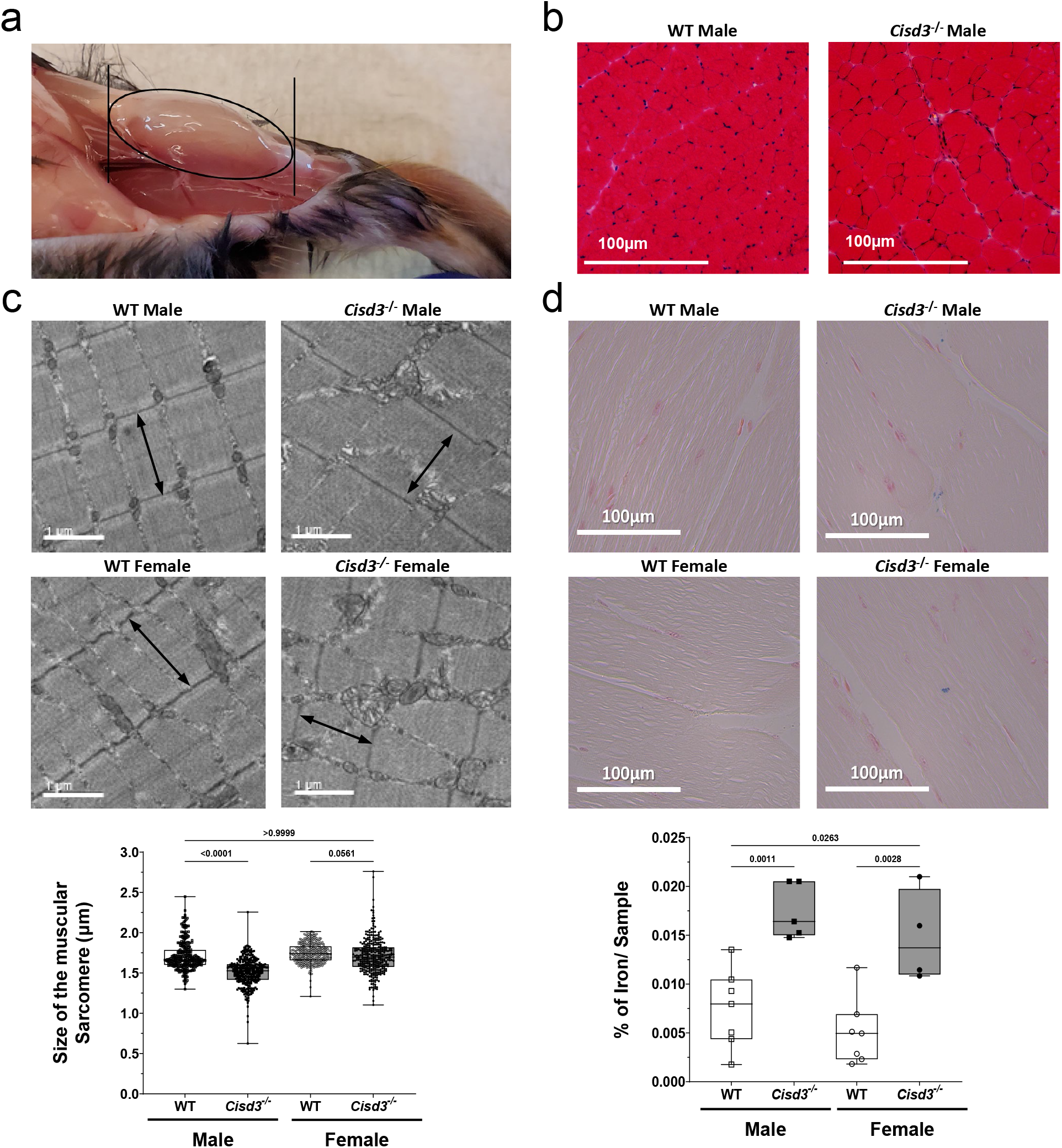
Skeletal muscles of *Cisd3^-/-^* mice display altered muscular sarcomere structure and accumulate iron. **a** A representative picture of the quadricep muscle of mice used for the comparative analysis of wild type (WT) and *Cisd3^-/-^* skeletal muscles. **b** Representative images of cross sections of WT and *Cisd3^-/-^* mice quadricep muscle stained with H&E (Fig. S2). **c** Representative transmission electron microscope (TEM) images of WT and *Cisd3^-/-^* mice quadricep muscle (top) and bar graphs showing quantification of images for sarcomere size (distance between Z-lines; bottom). **d** Representative Perls’ Prussian blue iron staining images of WT and *Cisd3^-/-^* mice quadricep muscle (top) and bar graphs showing quantification of images for muscle iron content (bottom). Results are shown for male and female separately and presented as mean ± SD of 6 different animals (3 different males and 3 different females) from each group (WT and *Cisd3^-/-^* mice). Two-way ANOVA followed by a Tukey test was used to calculate statistical significance. White box and white square, WT male mice; Gray box and black square, *Cisd3^-/-^* male mice; white box and white circle, WT female mice; Gray box and black circle, *Cisd3^-/-^* female mice. Abbreviations: CISD, CDGSH Iron Sulfur Domain; H&E, Hematoxylin & Eosin; KO, knock out; TEM, transmission electron microscope; WT, wild type.

As the deficiency in the level of CISD proteins correlates with enhanced levels of iron in cells^26,38^, we used the Perls’ Prussian blue iron stain method to measure the level of iron in muscle sections from WT and *Cisd3^-/-^* mice. This analysis revealed that muscle tissue from male and female *Cisd3^-/-^* mice contained higher levels of iron compared to male and female muscle sections from WT mice (Fig. 3d).

### Deficiency in CISD3 affects the structure and function of skeletal muscle mitochondria

Previous studies demonstrated that deficiency in CISD3 (as well as CISD1 or CISD2) results in mitochondrial structural abnormalities and increased mitochondrial damage in cancer cells^4,26,39^. We therefore studied the structure of mitochondria in male and female *Cisd3^-/-^* mice using TEM. This analysis revealed that, compared to male and female WT mice, mitochondria from male and female *Cisd3^-/-^* mice are abnormal and appear swollen with damaged crista (Fig. 4a).

**Fig. 4:**
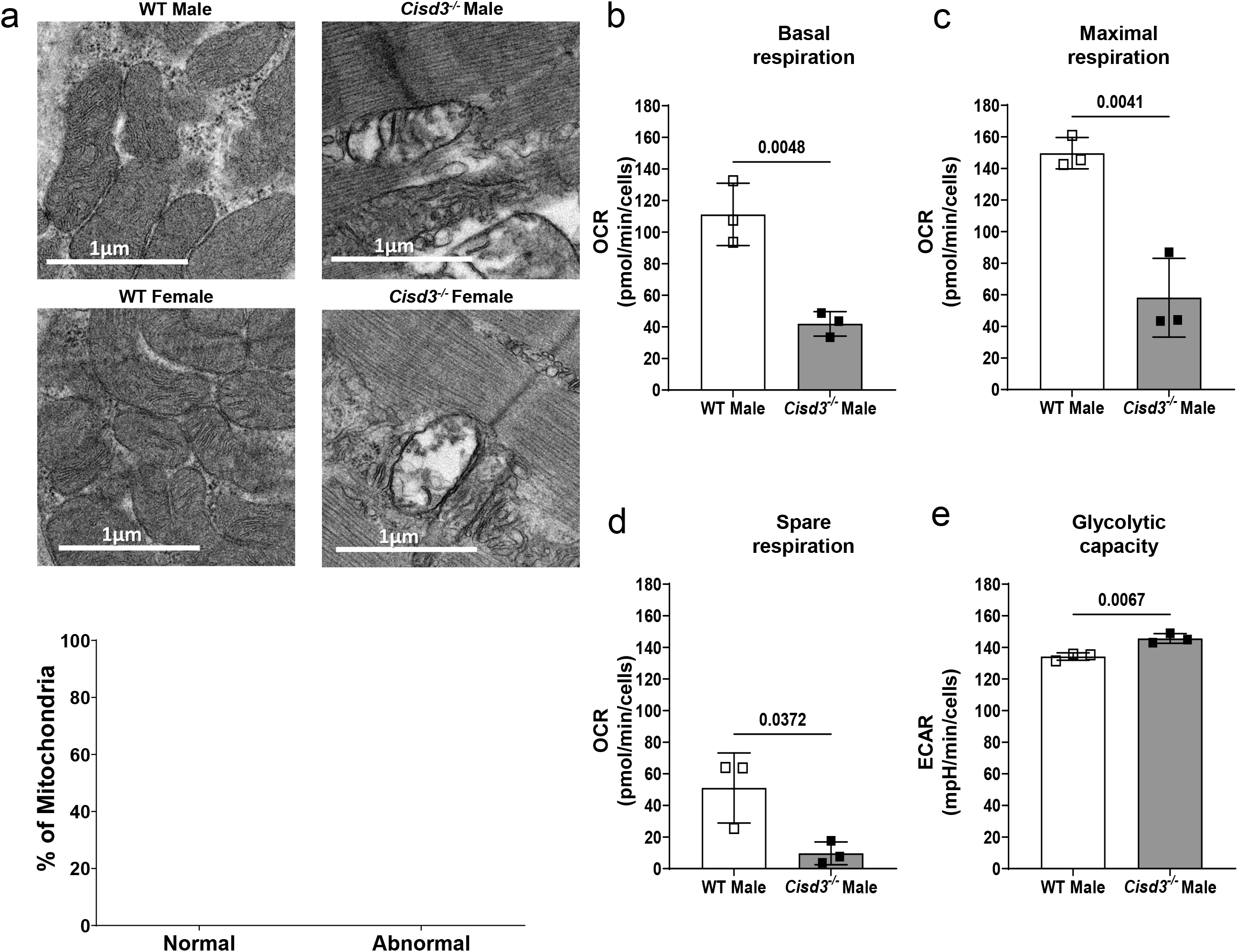
Impaired structure and function of mitochondria from skeletal muscles of *Cisd3^-/-^* mice. **a** Representative transmission electron microscope (TEM) images of mitochondria from wild type (WT) and *Cisd3^-/-^* mice (top) and bar graphs showing quantification of mitochondrial damage (bottom). **b-e** Bar graphs for basal (b), maximal (c), and spare (d) respiration, and glycolytic activity (e) of skeletal muscle fibers from WT and *Cisd3^-/-^* mice (measured with an XFe24 Seahorse apparatus). Results are shown for male mice and presented as mean ± SD of 6 different animals from each group (WT and *Cisd3^-/-^* mice). Two-way ANOVA followed by a Tukey test was used to calculate statistical significance. White box and white square, WT male mice; Gray box and black square, *Cisd3^-/-^* male mice; white box and white circle, WT female mice; Gray box and black circle, *Cisd3^-/-^* female mice. Abbreviations: CISD, CDGSH Iron Sulfur Domain; ECAR, Extracellular Acidification Rate; KO, knock out; OCR, Oxygen Consumption Rate; TEM, transmission electron microscope; WT, wild type.

To test whether the damage to mitochondria in *Cisd3^-/-^* mice resulted in suppressed respiration, we used an XFe24 Seahorse apparatus to measure respiration and glycolysis in muscle fibers from WT and *Cisd3^-/-^* mice. As the phenotype of the CISD3 deletion appeared to be more severe in male mice (*e.g.,* Figs 2a, S2), we focused our studies henceforth on male mice. In agreement with the analysis of mitochondrial structure (Fig. 4a), and compared to muscle fibers from WT male mice, the basal respiration, maximal respiration, and spare respiratory capacity of muscle fibers from *Cisd3^-/-^* mice was significantly reduced (Figs. 4b-4d). In contrast, the glycolytic capacity of muscle fibers from *Cisd3^-/-^* mice was higher than that of male WT fibers (Fig. 4e).

### Proteomics analysis of skeletal muscles from *Cisd3^-/-^* mice

To determine the molecular function of CISD3 in mice skeletal muscles, we conducted a comparative proteomics analysis of quadriceps muscle tissue from 44-47-week-old WT and *Cisd3^-/-^* male mice (Fig. 3). Our analysis identified a total of 2,545 proteins. Of these, 804 proteins (32%) had a significant change in expression (up or down) in *Cisd3^-/-^* mice compared to WT (Fig. 5a; Tables S1, S2). Of the 804 proteins, 650 proteins (26%) decreased in their expression, while the expression of 154 proteins (6%) was elevated (Fig. 5a; Tables S1, S2).

**Fig. 5:**
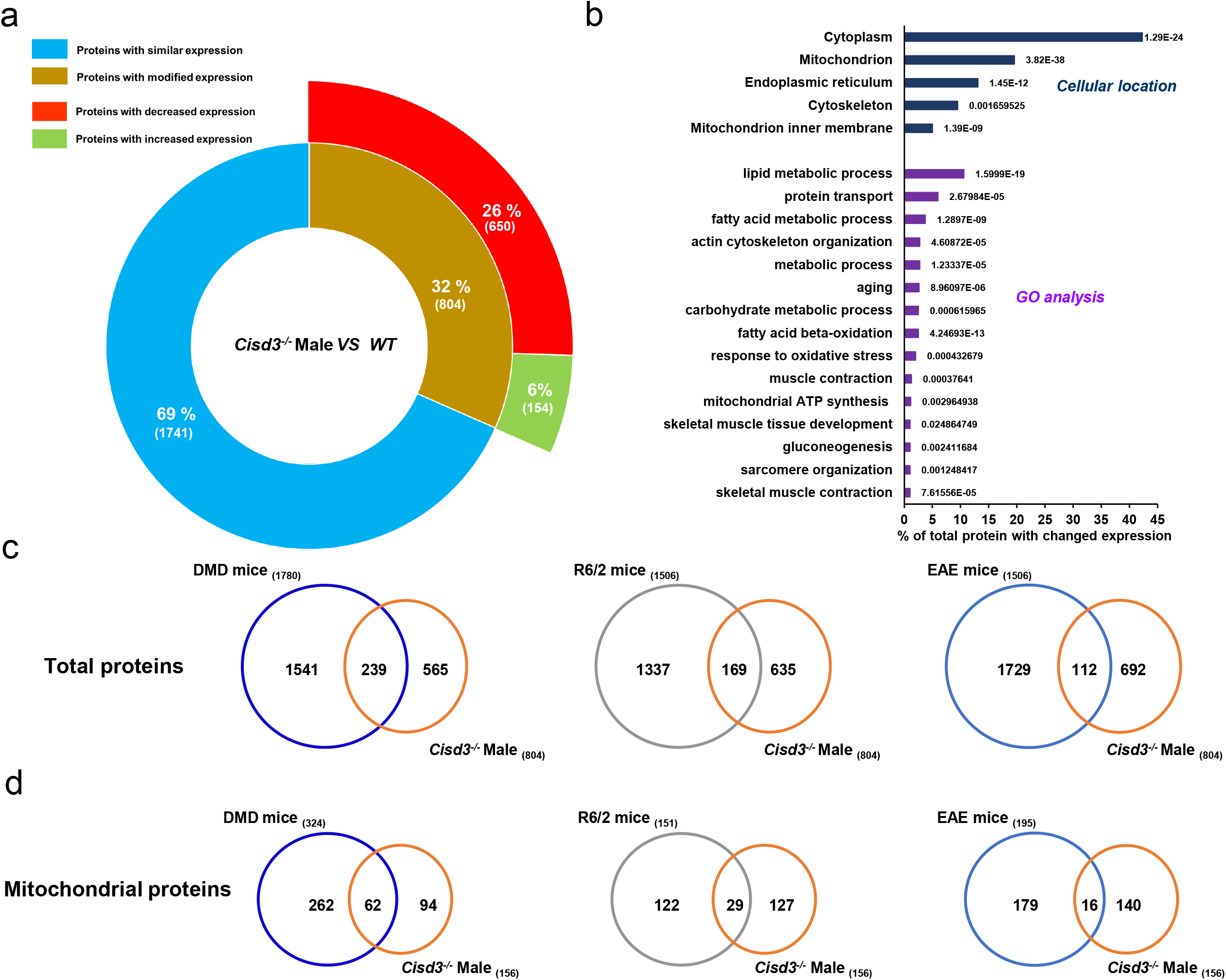
Proteomics analysis of skeletal muscles from wild type and *Cisd3^-/-^* mice. **a** A diagram showing the number of proteins altered in *Cisd3^-/-^* mice compared to wild type (WT) in quadricep muscle tissues of 44 week-old mice. **b** Subcellular localization and gene ontology (GO) annotation of proteins altered in *Cisd3^-/-^* mice compared to WT. **c** Venn diagrams showing the overlap between proteins altered in *Cisd3^-/-^*mice and proteins altered in mice model systems for Duchenne Muscular Dystrophy (DMD), Huntington Disease (HD; R6/2), and Multiple Sclerosis (MS; EAE). **d** Same as in c but for mitochondrial proteins. Results are shown for male mice and presented as mean ± SD of 6 different animals from each group (WT and *Cisd3^-/-^*mice). Two-way ANOVA followed by a Tukey test was used to calculate statistical significance. Abbreviations: CISD, CDGSH Iron Sulfur Domain; DMD, Duchenne muscular dystrophy; EAE, experimental allergic encephalomyelitis; HD, Huntington disease; GO, gene ontology; KO, knock out; MS, multiple sclerosis; WT, wild type.

Annotation of subcellular localization of the 804 proteins (significantly altered in their expression in *Cisd3^-/-^* mice compared to WT) revealed that most of these proteins localized to the cytosol and mitochondria (Fig. 5b). Gene ontology (GO) analysis of the 804 proteins identified lipid and carbohydrate metabolism, cytoskeleton, oxidative stress, aging, hypoxia, iron homeostasis, and gluconeogenesis among the most abundant proteins in this group (Fig. 5b). These findings are in agreement with our analysis of mitochondrial structure and function, supporting the observed reduction in respiration, coupled with the enhanced rate of glycolysis in *Cisd3^-/-^* mice (Figs. 4b-4e), as well as the overall damage to mitochondria (Fig. 4a).

To determine the degree of similarity between the proteomic changes occurring in *Cisd3^-/-^* mice and changes in protein expression occurring in mice models of different diseases, associated with muscle atrophy/ dystrophy, we compared the changes in the proteome of *Cisd3^-/-^*mice to that of DMD^40^, Huntington disease^41^, and multiple sclerosis (MS) ^42^. Interestingly, over 29% of proteins altered in *Cisd3^-/-^* mice were also altered in DMD mice, while only 21 and 14% of proteins altered in *Cisd3^-/-^* mice were also altered in Huntington diseases and MS mice, respectively (Fig. 5c). When the comparison of proteome similarity between *Cisd3^-/-^*, DMD, Huntington, and MS mice was restricted to mitochondrial proteins, the overlap between *Cisd3^-/-^* mice, DMD, Huntington, and MS was 66, 22, and 11%, respectively (Fig. 5d). The high overlap between *Cisd3^-/-^* and DMD (at the total and mitochondrial protein level; Figs. 5c, 5d) could suggest that CISD3 and DYSTROPHIN deficiency share some similarities with respect to their effect(s) on mitochondrial function and proteome alterations. In that respect, it is interesting to note that the abundance of several proteins involved in the DYSTROPHIN muscle complex (NOS1, DAG1, and Sgcd/Delta-sarcoglycan), was significantly reduced in *Cisd3^-/-^* male mice (Fig. S3; DYSTROPHIN itself was not significantly altered in *Cisd3^-/-^*mice).

### Changes in the abundance of proteins associated with metabolic pathways in *Cisd3^-/-^* mice

To further investigate the proteomic changes associated with CISD3 deficiency, we examined changes in protein expression associated with different metabolic pathways (Fig. 6). This analysis revealed that the expression of many key proteins involved in the TCA cycle is suppressed in *Cisd3^-/-^* mice (Fig. 6a). The expression of several proteins involved in fatty acid oxidation, supplying Acetyl-CoA to the TCA pathway, was also suppressed (Fig. 6b). In addition, and in agreement with the enhanced glycolytic activity of muscle fibers from *Cisd3^-/-^* mice (Fig. 4e), the expression of several proteins involved in glycolysis was enhanced, while the expression of several proteins involved in gluconeogenesis (reversal of glycolysis) was suppressed (Fig. 6c). The findings presented in Fig. 6 suggest that CISD3 deficiency causes major alterations in mitochondrial metabolism and function that could result in the suppression of respiration (also supported by the reduced respiratory activity displayed by muscle fibers from *Cisd3^-/-^* mice; Figs. 4b-4d). We therefore focused our attention on the abundance of proteins involved in the mitochondrial respiratory electron transfer chain.

**Fig. 6:**
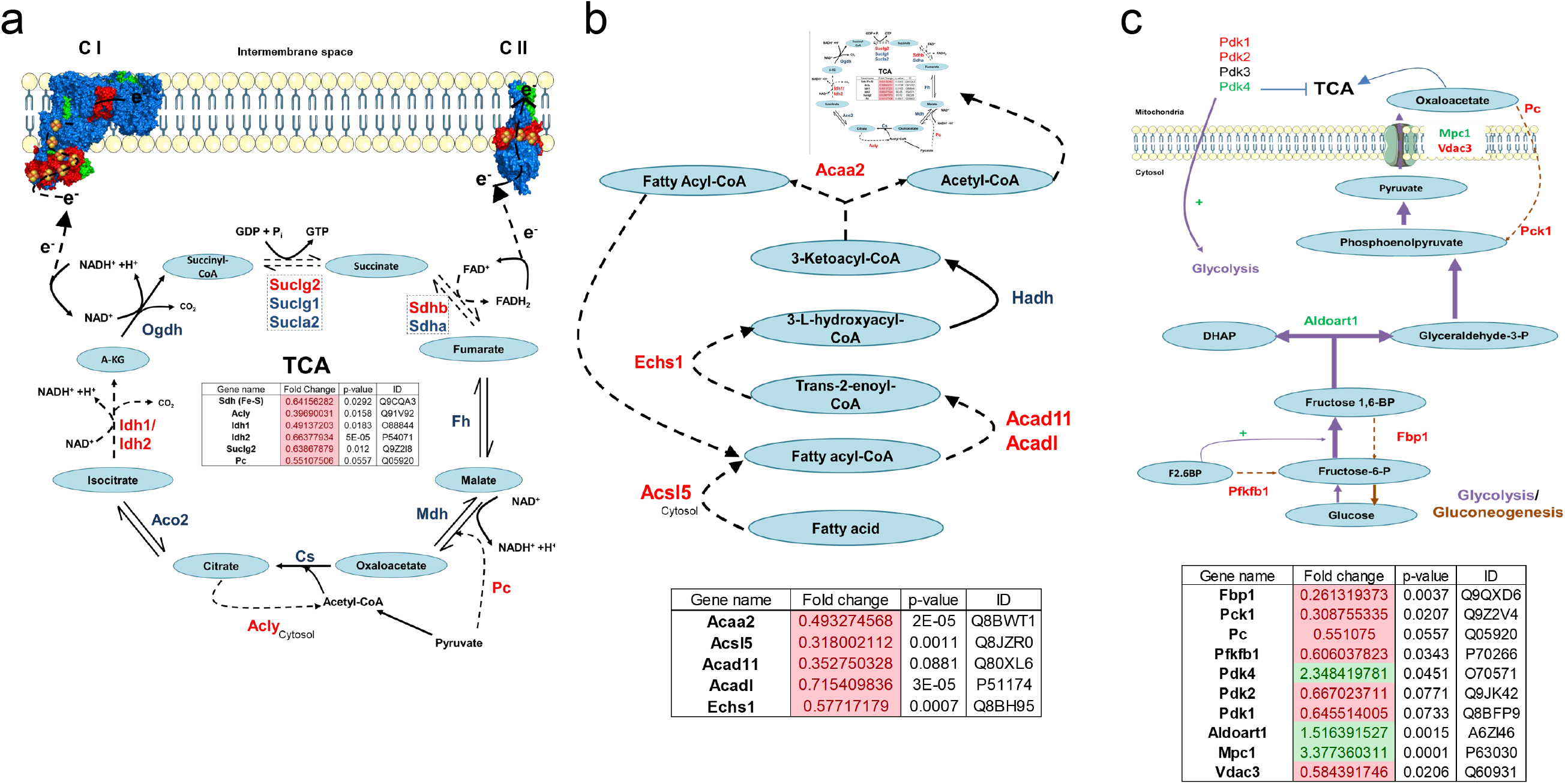
Changes in the abundance of proteins associated with metabolic pathways in *Cisd3^-/-^*mice. Changes in protein abundance in *Cisd3^-/-^* mice are shown for the tricarboxylic acid cycle (TCA; **a**), fatty acid oxidation (b), and glycolysis/gluconeogenesis (**c**) pathways. Results are shown for male mice. Abbreviations: Cs, citrate synthase; Acly, ATP-citrate synthase; Aco2, Aconitase 2; Idh, Isocitrate dehydrogenase [NADP]; Ogdh, Oxoglutarate Dehydrogenase; Suclg, Succinate--CoA ligase; Pc, pyruvate carboxylase; Sdh, Succinate dehydrogenase [ubiquinone]; Fh, Fumarate hydratase; Mdh, Malate dehydrogenase; A-KG, α-Ketoglutarate; Acaa2, 3-ketoacyl-CoA thiolase; Acsl5, Long-chain-fatty-acid— CoA ligase 5; Acad11, Acyl-CoA dehydrogenase family member 11; Acadl, Long-chain specific acyl-CoA dehydrogenase; Echs1, Enoyl-CoA hydratase; Fbp1, Fructose-1,6-bisphosphatase 1; Pck, Phosphoenolpyruvate carboxykinase; Pfkfb1, 6-phosphofructo-2-kinase/fructose-2,6-bisphosphatase 1; Pdk, Pyruvate dehydrogenase (acetyl-transferring)] kinase; Aldoart1, Fructose-bisphosphate aldolase; Mpc, Mitochondrial pyruvate carrier; Vdac3, Voltage-dependent anion-selective channel protein 3, F2.6BP, Fructose 2,6-bisphosphate; DHAP, Dihydroxyacetone phosphate.

Comparative analysis of Complex I, II, III, IV, and V protein levels between WT and *Cisd3^-/-^* mice revealed that CISD3 deficiency resulted in a rebalancing process between different subunits of these complexes (Fig. 7a). Expression of several proteins, including key [Fe-S] proteins in Complex I and II, was reduced, while expression of other proteins associated with these complexes was enhanced (Fig. 7a). Considering the reduction in the level of key proteins associated with the TCA cycle and Acetyl-CoA metabolism (Figs. 6a, 6b), involved in electron transfer to Complex I and II, in *Cisd3^-/-^* mice (Fig. 6), and the overall reduction in respiration (Figs. 4b-4d), this finding suggests that CISD3 could play an important role in maintaining proper metabolic reactions associated with respiration in muscles. Bearing in mind the soluble nature of CISD3^4^, its potential to transfer clusters to acceptor proteins^4,22^, and the fact that many of the proteins altered in their expression in Complex I were localized to the mitochondrial matrix-exposed part of this complex (Fig. 7a), we hypothesized that CISD3 is involved in donating [2Fe-2S] clusters/electrons to a subunit(s) of this complex (*e.g.,* the [2Fe-2S] subunit of Complex I, NDUFV2).

**Fig. 7:**
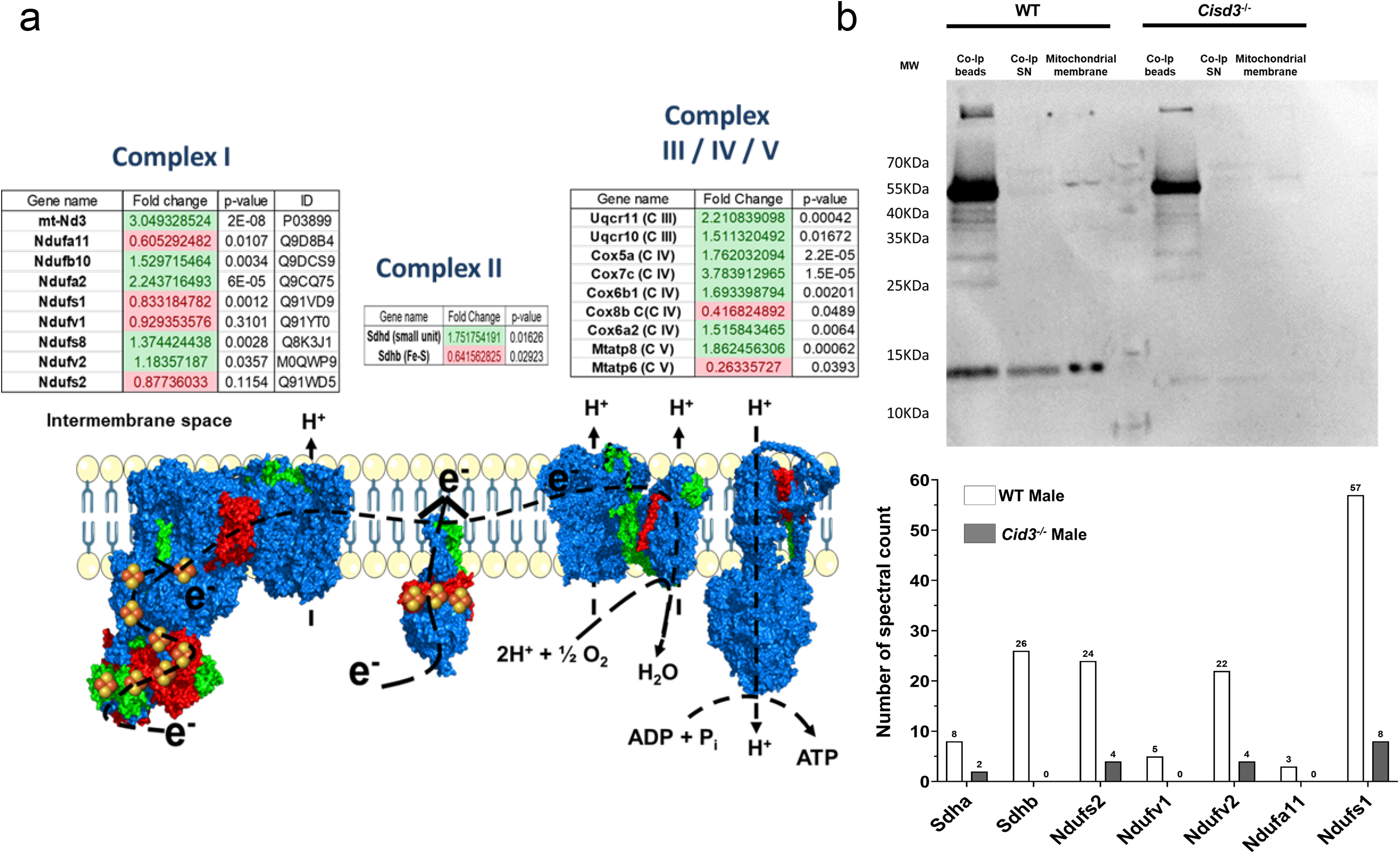
Abundance of proteins involved in the mitochondrial respiratory chain and identification of proteins associated with CISD3 in skeletal muscles. **a** Changes in protein abundance in *Cisd3^-/-^* mice are shown for complex I, II, and III/IV/V proteins. **b** Identification of proteins that associate with CISD3 *in vivo* in skeletal muscles following immunoprecipitation assays performed with skeletal muscles from wild type (WT) and *Cisd3^-/-^* mice. Protein identification was performed by proteomics analysis. Results are shown for male mice. Abbreviations: mt-Nd3, NADH-ubiquinone oxidoreductase chain 3; Ndufa, NADH dehydrogenase [ubiquinone] 1 alpha subcomplex subunit; Ndufb10, NADH dehydrogenase [ubiquinone] 1 beta subcomplex subunit 10; Ndufs, NADH:ubiquinone oxidoreductase core subunit; Ndufv, NADH:ubiquinone oxidoreductase core subunit V; Sdh, Succinate dehydrogenase [ubiquinone]; Uqcr, Cytochrome b-c1 complex subunit; Cox, Cytochrome c oxidase subunit 5A; Mtatp, ATP synthetase protein; Co-ip, Co-Immunoprecipitation; SN, Supernatant; MW, Molecular Weight; CISD, CDGSH Iron Sulfur Domain; KO, knock out; WT, wild type.

### Identification of proteins associated with CISD3 in mice skeletal muscles

To identify the different proteins that interact via protein-protein interactions with CISD3 in mice skeletal muscles, we conducted an immunoprecipitation study using quadriceps muscle tissues from WT and *Cisd3^-/-^* mice and an antibody against CISD3 (Fig. 1c)^4,22^. This analysis revealed that several proteins that are part of Complex I (NDUFS2, NDUFV1, NDUFV2, NDUFA11, and NDUFS1) and Complex II (SDHA and SDHB) associate with CISD3 *in vivo* in quadriceps muscles of WT (but not *Cisd3^-/-^* mice; Fig. 7b). Of these, the levels of NDUFA11, NDUFS1, and SDHB were reduced in their expression in *Cisd3^-/-^* mice (Fig. 7a). In addition, among the proteins identified as interacting with CISD3 *in vivo*, at least two contained [2Fe-2S] clusters: NDUFV2 and SDHB, of which NDUFV2 was part of the matrix-exposed part of Complex I (Fig. 7a). We therefore focused on the potential interaction of CISD3 with NDUFV2.

### *In vitro* interaction and cluster transfer between CISD3 and NDUFV2

To further study the potential interaction between CISD3 and NDUFV2 (Fig. 7b), we cloned the cDNAs of CISD3 and NDUFV2, overexpressed these proteins in bacteria, and purified them for *in vitro* studies. Co-incubation of purified holo-CISD3 and holo-NDUFV2 proteins, followed by immunoprecipitation and proteomics analysis, revealed that the two proteins physically interact *in vitro* (Figs. 8a-8c). As β-mercaptoethanol prevented this interaction (Figs. 8a, 8b), it is possible that it involves the formation of disulfide bridges within one or both proteins and/or between the two different proteins. To examine whether the physical interaction between CISD3 and NDUFV2 (Figs. 7b, 8a-8c) results in a cluster-transfer reaction (between CISD3 and NDUFV2), we studied the potential of CISD3 to transfer its [2Fe-2S] clusters to NDUFV2. Co-incubation of purified holo-CISD3 with purified apo-NDUFV2 resulted in the transfer of the [2Fe-2S] clusters of CISD3 to NDUFV2 (evident by the shift in the spectrum of apo-to holo-NDUFV2; Fig. 8d). For a cluster transfer reaction to occur, the distance between the two cluster binding sites of CISD3 and NDUFV2 needs to be in the 10-15 Å range^25^. To determine if the interaction of CISD3 with NDUFV2 brings the cluster binding domains of the two proteins within this range, we conducted a computer-simulated binding study of the two proteins. The computer modeled binding structure of CISD3 and NDUFV2 revealed that the two proteins interact (driven by electrostatic and hydrophobic interactions) and that their clusters are about 12.5 Å apart when they interact (Fig. 8e). This distance would allow cluster transfer from CISD3 to NDUFV2.

**Fig. 8:**
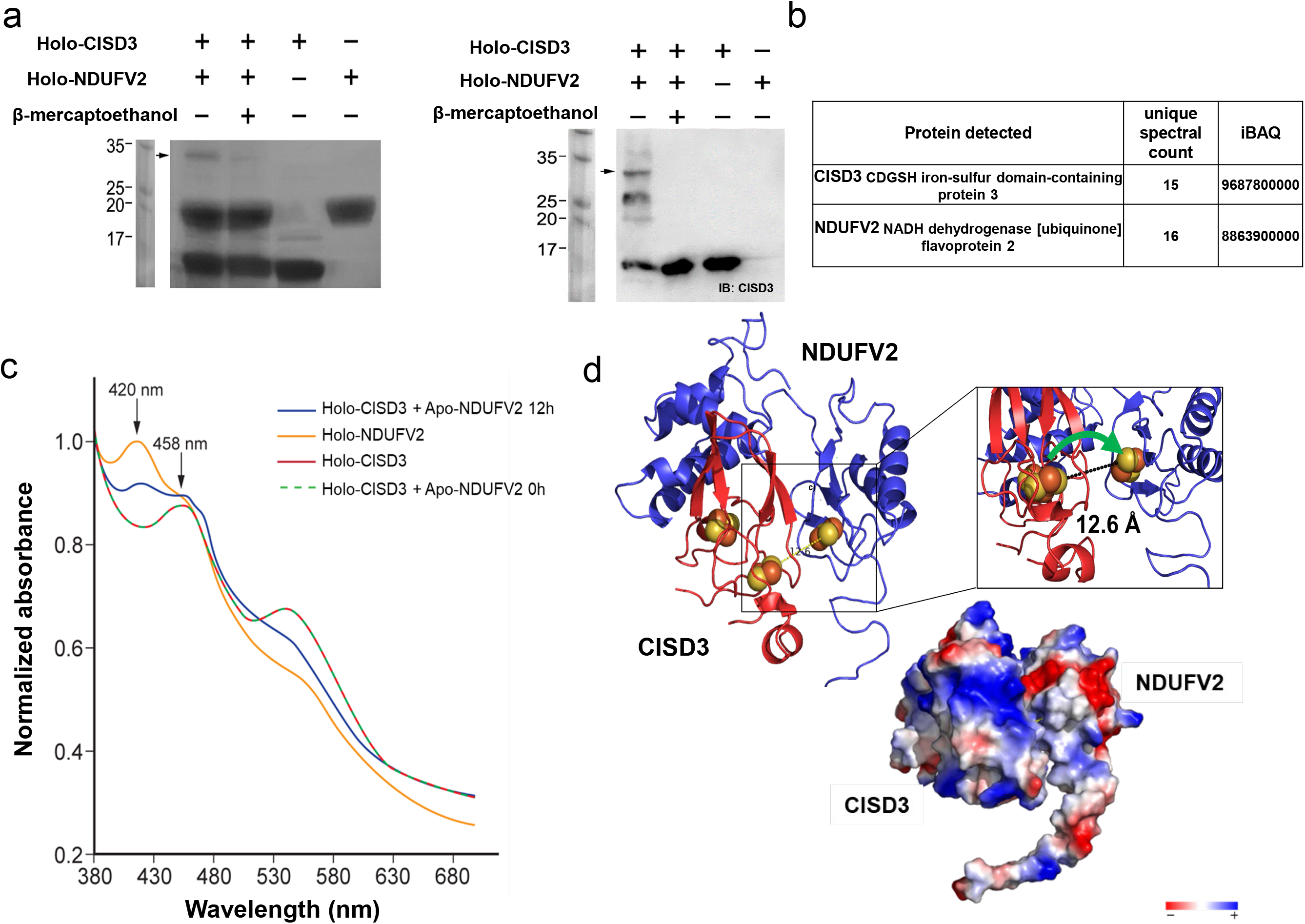
*In vitro* protein-protein interaction and cluster transfer between CISD3 and NDUFV2. **a** Protein gel (left) and protein blot with antibody against CISD3 (right) showing *in vitro* protein-protein interaction between CISD3 and NDUFV2. Arrows indicate the location of the protein-protein complex between CISD3 and NDUFV2. **b** Identification of proteins found in the band indicated by arrows in (a) using proteomics analysis. **c** Cluster transfer from holo-CISD3 to apo-NDUFV2 visualized by changes in spectra of the two proteins following co-incubation. **d** Computer-optimized model for the protein-protein interaction between CISD3 and NDUFV2. The distance between the [2Fe-2S] clusters of the proteins is shown to be 12.6Å. Abbreviations: CISD, CDGSH Iron Sulfur Domain; Ndufv2, NADH:ubiquinone oxidoreductase core subunit V2; IB, Immunoblotting; IBAQ, intensity-based absolute quantification.

## DISCUSSION

CISD3 was proposed to function as part of an iron-sulfur cluster relay mechanism, transferring [2Fe-2S] clusters from within the mitochondria to CISD1 (localized on the OMM), and from there, *via* CISD2, to the cytosolic iron sulfur cluster (ISC) biogenesis complex of the cytosol (*via* Anamorsin and perhaps other proteins)^2,22,24,25^. However, very little is currently known about the function of CISD3 within the mitochondria. In addition, most studies of CISD3 function were focused on its role in regulating ferroptotic and apoptotic cell death in cancer cells and *C. elegans*^4,28,31,32,43^. In contrast, very little is known about the physiological role of CISD3 in mammalian cells and tissues. Here we show that CISD3 plays a key role in the maintenance and function of skeletal muscles, supporting respiration and other mitochondrial metabolic pathways *via* interacting with different subunits of Complex I and II (Figs. 2-8). In particular, we show that CISD3 can physically interact with and transfer its clusters to NDUFV2, an [2Fe-2S]-containing protein that is localized to the mitochondrial matrix-exposed part of Complex I (Figs. 8, 9). Deficiency in CISD3 therefore causes reduced respiration, enhanced glycolysis, mitochondrial structural damage, and muscle atrophy (Figs. 2, 4).

In cancer cells all three CISD proteins were shown to function in a similar role, regulating mitochondrial iron and ROS levels and suppressing apoptotic and ferroptotic cell death^2,3,26,44^. As described above, they were also proposed to function as part of a [2Fe-2S] relay that transfers Fe-S clusters from within the mitochondria to the cytosol^22^. In contrast, while all three CISD proteins were found to play a role in muscle maintenance and function^11,12,38, 45–47^ (Figs. 2-7), they appear to have different functions in muscle tissues. CISD2 was shown to regulate calcium signaling and autophagy in skeletal muscles, as well as to protect muscle cells from overaccumulation of iron^12,38,48^, CISD1 was shown to protect heart muscle cells and tissues from iron and ROS accumulation, and ferroptosis activation^11,49^, and the present study reveals that CISD3 plays a key role in supporting the mitochondrial respiratory electron transport chain by providing [2Fe-2S] cluster to NDUVF2 (Figs. 4, 7-9). This new role for CISD3 demonstrate that under physiological conditions CISD proteins may mediate a variety of cellular roles that differ depending on their subcellular localization. While CISD2 that is localized to the OMM, ER, and MAM, is involved in regulating apoptosis, ferroptosis, calcium signaling, and autophagy *via* interactions with key signaling proteins found associated with these different membrane systems, CISD1 that is localized on the OMM, and CISD3 that is localized inside the mitochondria, are primarily involved in regulating the flow of [2Fe-2S] clusters from the mitochondria to the cytosol, thereby controlling mitochondrial iron and ROS levels and suppressing ferroptosis. In addition, our study reveals that CISD3 support respiration from within the mitochondria by providing [2Fe-2S] clusters to NDUFV2 (Figs. 4, 7-9).

NDUFV2 encodes a 27-kDa subunit of the Complex I N module^50,51^. Mutations in NDUFV2 were reported to cause hypertrophic cardiomyopathy and encephalopathy, as well as Leigh syndrome, a severe neurological and respiratory disorder that typically results in mortality (due to respiratory failure) at ages 2-3^52–54^. Our findings that CISD3 physically interacts with, and transfers its clusters to, NDUFV2 (Fig. 8), and is important for supporting overall respiration and mitochondrial integrity (Fig. 4), could implicate CISD3 in some of the different conditions described above, associated with NDUFV2 deficiency, as well as in additional mitochondrial complex I deficiencies, such as fatal neonatal lactic acidosis, leukoencephalopathy, hepatopathy, cardiomyopathy, childhood-onset mitochondrial encephalomyopathy, and stroke-like episodes^54–56^. The potential involvement of CISD3 in these and other presentations associated with mitochondrial/Complex I function should be explored in future studies, as CISD3 could become a molecular marker for some of these conditions.

A common thread that appears to be shared by all three CISD proteins is their involvement in supporting enhanced metabolic activity of cells. This theme is evident in the important role CISD proteins play in disorders such as diabetes, cancer, and neurodegeneration, as well as in their role in muscle tissues and the germline of *C. elegans*^1–3,57,58^. Interestingly, among the three CISD proteins, CISD3 is the most ancient and can be found in archaea and bacteria, as well as different eukaryotic cells and animals^33,34,59^. Our findings that CISD3 plays an important role in supporting respiration in muscle tissues could shed new light on the evolution of this protein and the role it played in the origin of the more ‘modern’ CISD proteins. Once mitochondria became a part of eukaryotic cells, the function of CISD3 as a cluster donor to respiratory complexes could have been expanded *via* gene duplications to CISD1 and CISD2^34^, to support [2Fe-2S] metabolism in the cytosol, and CISD1/2 acquired additional cytosolic roles related to coordinating mitochondrial-cytosolic interactions. The fact that CISD3 is generally absent in plants^33,34,59^, could suggest that once the chloroplast was acquired by eukaryotic cells, a different protein could have overtaken the role of CISD3, or that CISD1/2 that were already present in these early cells became the potential progenitors of the plant CISD proteins, as exemplified by the *Arabidopsis thaliana* AtNEET protein^8,60,61^. The fact that our *Cisd3^-/-^* mice survived to 60 weeks strongly suggest that additional molecular mechanisms can support, or at least partially replace, the function of CISD3 in mammalian cells (perhaps similar to the mechanisms replacing the function of this protein in plants). However, once the energetic demand from cells becomes high (as in the skeletal muscles of *Cisd3^-/-^* mice, or in cancer cells), over time, these alternative mechanisms may not be sufficient to replace CISD3 function, and mitochondrial integrity and function is hindered. In future studies the identity of the alternative mechanisms that replace CISD3 function should be addressed as they may be highly important for manipulating the enhanced metabolic state of cells in the presence or absence of CISD3.

The similarity in proteomic patterns between *Cisd3^-/-^* mice and a mice model for DMD (Fig. 5) is intriguing. Duchenne and Becker muscular dystrophies (DMD and BMD) are caused by a mutation in DYSTROPHIN, a protein that links cytoskeletal F-actin with the extracellular matrix^62–65^. Muscles without DYSTROPHIN are more sensitive to damage, resulting in progressive loss of muscle tissue and function^66^. It is possible that the similarity between *Cisd3^-/-^* and DMD mice is due to the enhanced muscle damage in *Cisd3^-/-^* mice (a result of mitochondrial dysfunction and structural impairment; Fig. 4). Alternatively, it could be due to the decrease in DYSTROPHIN complex proteins in *Cisd3^-/-^* mice (Fig. S3). It is also possible that at least part of the muscle damage in DMD mice is caused by respiratory complex failure, that is partially similar to the phenotype observed in *Cisd3^-/-^* mice (Fig. 4). Production and analysis of double *Cisd3*^-/-*/*^*Dystrophin*^-/-^ mice may help unravel this question. Overall, the similarity between DMD and *Cisd3^-/-^*model mice support a role for mitochondrial respiration impairment in DMD and could highlight new avenues for the proposed gene therapy of DMD^66,67^.

Several different drugs were shown to stabilize or destabilize the clusters of CISD1 and CISD2 proteins, including pioglitazone, used in the treatment of diabetes^28,68–71^. Due to the high similarity in structure between the different CISD proteins^1,3^, and considering the new findings revealed by this work regarding the role of CISD3 in muscle function and maintenances (Fig. 9), it might be interesting to test the effect of some of these drugs on muscle performance in control mice. Such studies could be important in aging mice, as the levels of the different CISD proteins in muscle tissues declines with age^15,45,46,48,57,58^. Specific drugs could also be developed to target CISD3, and these could impact muscle degeneration syndromes, aging, and different sport applications. Taken together, our study reveals a distinct physiological role for CISD3 as a cluster doner for mitochondrial proteins essential for supporting respiration and other metabolic activities in skeletal muscles.

**Fig. 9:**
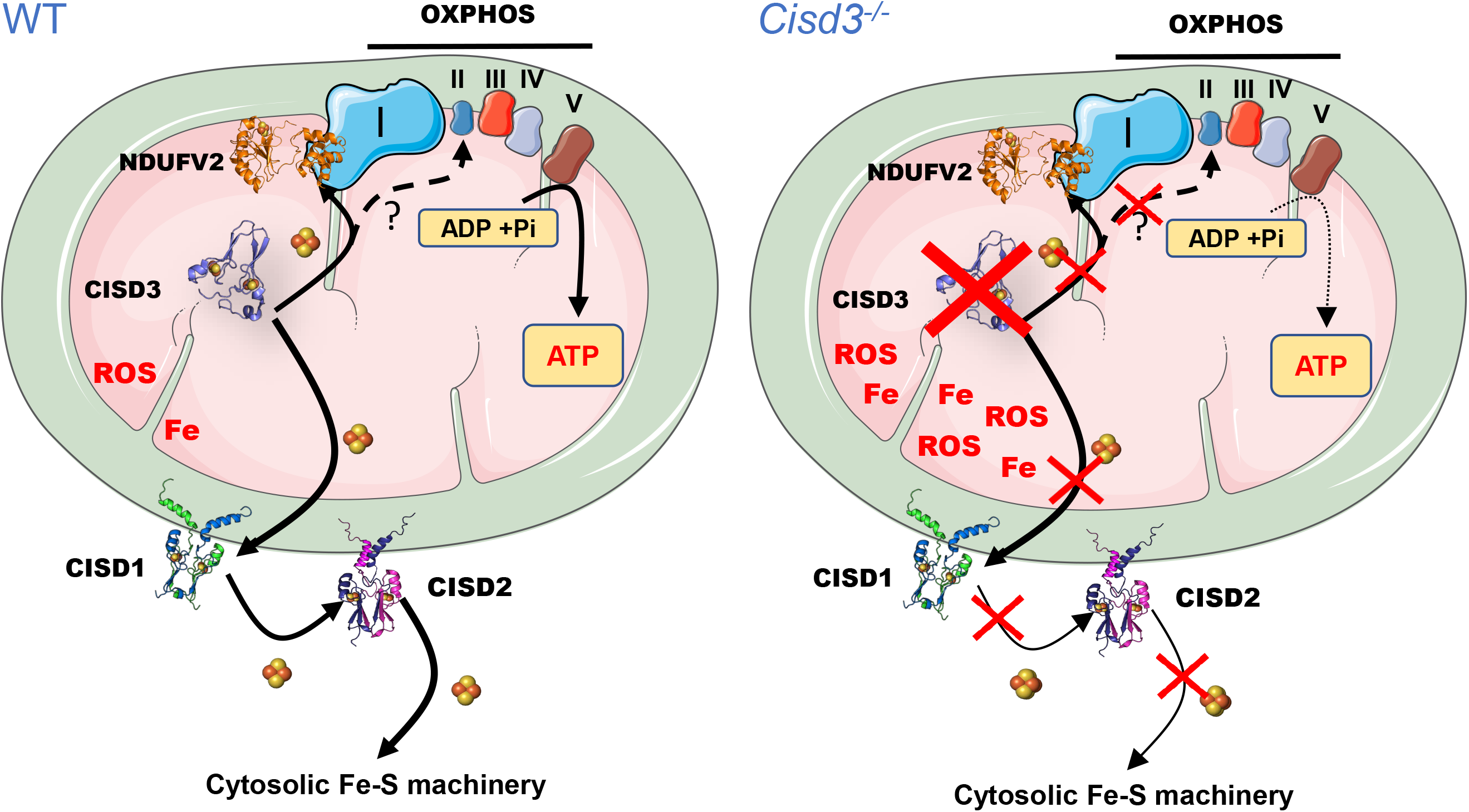
A model for the function of CISD3 in mitochondria from skeletal muscles of mice. CISD3 is shown to donate its [2Fe-2S] clusters to NDUFV2 (and perhaps other 2Fe-2S proteins of complex I and II) thereby supporting the function of the mitochondrial respiratory chain in muscle mitochondria and enable enhanced metabolic activity. Based on previous studies in cancer cells, CISD3 is also proposed to function as a part of a [2Fe-2S] cluster relay that transfer clusters from within the mitochondria to the cytosol via interactions of cISD3 with CISD1 and CISD3. In the absence of CISD3 the function of the mitochondrial respiratory chain is reduced, and mitochondria are more prone to iron overload and structural damage.

## Material and Methods

### Production of *Cisd3^-/-^* mice

All animals used in this study were housed under pathogen-free barrier conditions in an animal care facility and received humane care in compliance with the Principles of Laboratory Animal Care formulated by the National Society of Medical Research and the Guide for the Care and Use of Laboratory Animals (https://www.ncbi.nlm.nih.gov/books/NBK54050/). All experimental procedures were approved by the Institutional Animal Care and Use Committee of the University of Missouri (https://research.missouri.edu/acqa/acuc). C57BL/6NTac-Cisd3^em1(floxed^ ^ex2)Davis^ heterozygous mice were generated by the University of Missouri Animal Modelling Core (https://research.missouri.edu/animal-modeling), and bred to homozygous status. C57BL/6NTac-Cisd3^em1(floxed^ ^ex2)Davis^ mice were then bred to B6.C-Tg(CMV-cre)1Cgn/J mice from Jackson Laboratory (stock 006054)^72^, and homozygous *Cisd3^-/-^*mice were generated (Fig. 1a, 1b). The following primer sequences were used for genotyping the mice (Fig. 1a, 1b): forward primer 5’-AGGACAGCCACCTGATTTCAAGC-3’, reverse primer 5’-CTAACCGTTAGCGTCAGAGATCAGG-3’, which produced a wild-type band of 304 bp and a mutant band of 162 bp (Fig. 1a, 1b).

### *Cisd3^-/-^* mice phenotyping

The physical condition, body weight, and general condition of the animals were recorded on a weekly basis, until the day of the termination. Weight was measured by placing the mice in an empty sterilized pipet tip container and data was recorded on Ohaus PS121 portable balance (Fig. S1). The grip strength of control and *Cisd3^-/-^* mice was measured using a homemade wheel which consisted of a metal Ball bearing roller on which a circular 2 mm thick loop was suspended on a support stand (25 cm of diameter at a height of 66 cm, Fig. S4a). Mice were allowed to hang from the bottom of the wheel and timed until they fell off into a soft container. A 2 mm thick straight brass rod was also held by a support stand horizontally and time until mice fell off from it into a soft container was determined (Fig. S4b). Male and female C57BL/6J wild type (Control) and *Cisd3^-/-^*mice were euthanized at the age of 47 weeks. Mice were placed under anesthesia using 4% Isoflurane, sacrificed, and subjected to Quadriceps tissue (Fig. 3a) collection.

### Protein blot analysis

Quadriceps tissue was collected, immediately flash-frozen in liquid nitrogen, and stored at −80 °C. Muscle tissues were lysed by grinding to a fine powder in liquid nitrogen and then resuspended in 20 mm Tris pH 8.0, 150 mm NaCl, 1% Triton 100-X, 10% glycerol, 5 mm EDTA, 5 ng mL^−1^ aprotinin, and 5 mm PMSF and kept on ice for 2 h with vortexing every 30 min. Following centrifugation at 20 000 g for 15 min, the supernatant was denatured with Laemmli sample buffer, as described in^38^. Protein content was determined by The Pierce 660 nm Protein Assay (catalog number 1861426). Following SDS-PAGE separation of equal amounts of proteins, proteins were transferred to nitrocellulose blots and incubated with antibodies (in-house produced) against CISD3, CISD2 and CISD^4,12, 22–27,38,39^, and β-actin (Abcam, MA). Peroxidase-conjugated Affinity Pure goat anti-rabbit IgG from Jackson ImmunoResearch Laboratories (Jackson ImmunoResearch; West Grove, PA) was used as secondary antibody.

### Hematoxylin and eosin (H&E) staining of muscle tissue

Quadriceps tissue was collected and snap frozen by immersing it in liquid 2-methyl butane. After a quick evaporation time, the frozen tissues were completely covered with optimal cutting temperature (OCT) embedding compound in a cryomold and then immersed in liquid nitrogen until completely frozen. Twelve-micron sections were prepared using a cryostat (Leica CM1520)) and subjected to H&E staining^73^ at the Histopathology Department of the Veterinary Medical Diagnostic Laboratory (University of Missouri; https://vmdl.missouri.edu/).

### Transmission electron microscopy (TEM) analysis of muscle tissue

Unless otherwise stated, all reagents were purchased from Electron Microscopy Sciences and all specimen preparation was performed at the Electron Microscopy Core Facility, University of Missouri (https://emc.missouri.edu/). Quadriceps tissue was collected, sliced into 1 mm sections, and fixed in 2% paraformaldehyde, 2% glutaraldehyde in 100 mM sodium cacodylate buffer pH 7.35. Tissue slices were then rinsed with 100 mM sodium cacodylate buffer, pH 7.35 containing 130 mM sucrose. Secondary fixation was performed using 1% osmium tetroxide (Ted Pella, Inc. Redding, California). Specimens were then incubated at 4 °C for 1 hr, rinsed with cacodylate buffer, followed by rinsing in with distilled water. En bloc staining was performed using 1% aqueous uranyl acetate (4 °C overnight). A graded dehydration series was performed using ethanol at 4°C, transitioned into acetone, and dehydrated tissues were then infiltrated with Epon resin for 24 hours at room temperature and polymerized at 60°C overnight. Sections were cut to a thickness of 75 nm using an ultramicrotome (Ultracut UCT, Leica Microsystems, Germany) and a diamond knife (Diatome, Hatfield PA). Images were acquired with a JEOL JEM 1400 transmission electron microscope (JEOL, Peabody, MA) at 80 kV using a Gatan Rio CMOS camera (Gatan, Inc, Pleasanton, CA). Results were evaluated from 50-100 different sections randomly selected, every cell was evaluated for its nuclear and general morphology in low magnification. At a higher magnification, averaged over 20 fields per cell, at least 100– 200 mitochondria were counted. Mitochondrial damage was expressed as the ratio of damaged organelle to total number of the organelle. Distance between the Z-lines representing the sarcomere size, were evaluated for each field (15-20 fields/ mouse).

### Perls’ Prussian blue staining of muscle tissues

Quadriceps tissues were fixed in 4% paraformaldehyde, processed for paraffin mounting, and sectioned (12 µm). Slides prepared from each sample were subjected to iron staining using the Mallory’s iron stain (Perls’ Prussian blue) method^74^. Slides were deparaffinized and rehydrated with distilled water, then potassium ferrocyanide/HCL solution was added for 10 min. Percent (%) of stained cells per field were calculated in the *Cisd3^-/-^ group* and compared to control. Data was acquired on an Invitrogen (Waltham, MA) EVOS XL Core Configured Cell Imager with Mechanical Stage. Results are shown for male and female and presented as mean ± SD for six different animals (three different males and three different females) from each group (control and *Cisd3^-/-^*), As described in^38^.

### Muscle fibers isolation

Full Quadriceps muscle tissues were collected, rinsed in PBS, and incubated in collagenase A (4 mg/mL) prepared in Dulbecco’s Modified Eagle Medium (D-MEM) high glucose, no sodium pyruvate or phenol red (Invitrogen, Waltham, MA, 21063-029), FBS (2 %), and collagenase A (4 mg/mL; Sigma-Aldrich, St. Louis, MO, COLLA-RO) pH 7.4 for 30 minutes. The fibers were separated using a wide bore p1000 pipette to yield single quadriceps myofibers in D-MEM Medium (Invitrogen, Waltham, MA, 21063-029), FBS (2 %). The separated myofibers were then plated in an Agilent (Santa Clara, CA) Seahorse XF24 Islet capture Microplates and covered with a grid to reduce the movement of the fibers and incubated for 1 hr in an incubator at 37°C and 5% CO2.

### Seahorse analysis of muscle fibers from control and Cisd3^-/-^ mice

For mitochondrial measurements: 1 hr before the experiment, the fibers were placed in an incubator at 37 °C with no CO2 and the medium was replaced with XF Base media (Agilent, Santa Clara, CA) with glucose (10 mM), sodium pyruvate (1 mM) and L-glutamine (2 mM) (Gibco, Waltham, MA), pH 7.4 at 37 °C. Fiber oxygen consumption rate (OCR) was measured during the Seahorse Mito Stress assay (Agilent; Santa Clara, CA), with the addition of oligomycin (0.65 µM), carbonyl cyanide 4-(trifluoromethoxy) phenylhydrazone (FCCP; 1.5 µM) and antimycin A and rotenone (1 μM each). Measured parameters were defined as follows: 3 min mix, 3 min wait, 3 min measurement, repeated 3–4 times at basal and after each addition. For glycolysis measurements, 1 hr before the analysis, the fibers were placed in an incubator at 37 °C with no CO2 and the medium was replaced with XF Base media (Agilent; Santa Clara, CA), with sodium pyruvate (1 mM) and L-glutamine (2 mM) (Gibco, Waltham, MA), and without glucose at pH 7.4 at 37 °C). Fiber extracellular acidification rate (ECAR) was measured during a Seahorse Glycolysis Stress assay (Agilent, Santa Clara, CA) with the addition of Glucose (10 mM), oligomycin (2 µM) and 2-Deoxy-D-glucose (2-DG: 50 µM). Measured parameters were defined as follows: 3 min mix, 3 min wait, 3 min measurement, repeated 3–4 times at basal and after each addition.

### Proteomics analysis

Quadriceps tissue was collected, immediately flash-frozen in liquid nitrogen, and stored at −80 °C. Muscle tissues were lysed by grinding to a fine powder in liquid nitrogen and resuspending in 20 mM Tris pH 8.0, 150 mm NaCl, 1% Triton 100-X, 10% glycerol, 5 mm EDTA, 5 ng mL−1 aprotinin, and 5 mM PMSF, and kept on ice for 2 hr with mixing every 30 min. Following centrifugation at 20 000 g for 15 min, the supernatant was denatured with Laemmli sample buffer, as described in^38^. The proteins were precipitated with cold acetone and stored at −20°C overnight. Protein pellets were recovered by centrifugation at 16 000 g for 10 min at 4°C and washed with 80% acetone. The final protein pellets were suspended in 6 M urea, 2 M thiourea and 100 mM ammonium bicarbonate. The protein concentration was determined using Pierce 660nm Protein Assay (Thermo-Fisher Scientific, Waltham, MA) according to the manufacturer’s protocol. 40 microgram proteins from each sample were reduced by 5 mM dithiothreitol (DTT) at room temperature for 1 hr and alkylated with 15 mM iodoacetamide (IAA) at room temperature for 30 min in the dark. The excess of IAA was quenched by adding 5 mM DTT and incubate for 15 min. Mass spectrometry-grade LyC (Promega, Madison, WI) was added at an enzyme-to-protein ratio of 1:50 (w/w) and incubated for 3 hr at 37 °C, and an additional mass spectrometry-grade trypsin was added at a 1:50 (w/w) ratio for digestion overnight at 37 °C. Digestion was quenched with 0.5% TFA. For experimental DDA library generation, 10 micrograms protein digests from each sample were pooled and loaded onto a high-pH RP fractionation spin column (Pierce, Waltham, MA) according to instructions. Eight fractionated peptide samples were dried and dissolved in 5% Acetonitrile/0.1% FA. The rest of the digested peptides were desalted using the C18 tip from Thermo Scientific (Waltham, MA). The desalted peptides were dried and resuspended in 5% Acetonitrile/0.1% FA. Peptide concentrations were measured optically at 280 nm (Nanodrop 2000; Thermo-Fisher, Waltham, MA), and 500 ng peptides were loaded onto a 20 cm long x 75 µm inner diameter pulled-needle fused-silica analytical column packed with Waters BEH-C18, 1.7um reversed phase resin (Waters, Milford, MA). Peptides were separated and eluted from the analytical column with a 90 min gradient of acetonitrile at 300 nL/min. The mobile phases comprised 0.1% FA as solution A and 0.1% FA/99.9% ACN as solution B. The Bruker nanoElute system (Bruker, Billerica, MA) was attached to a Bruker timsTOF-PRO mass spectrometer (Bruker, Billerica, MA) via a Bruker CaptiveSpray source (Bruker, Billerica, MA). The spectral library samples were acquired using the DDA-PASEF mode, with one MS frame and 10 PASEF/MSMS scans per topN acquisition cycle. Precursors with a charge of +1 were filtered out based on their position in the m/z-IM plane and only precursors with an intensity threshold of 2,500 arbitrary units were selected for fragmentation. Target MS intensity for MS was set at 10,000 arbitrary units, ion mobility coefficient (1/K0) value was set from 1.6 to 0.6 V cm-2, collision energy was set from 20-59 eV, MS data were collected over m/z range of 100 to 1700. Dynamic exclusion was activated after 0.4 minutes, and isolation width was set to 2 for m/z <700 or 3 for m/z >700. For data-independent acquisition, dia-PASEF method was used to cover an m/z range of 400 to 1200 and an ion mobility range of 0.57 to 1.47 Vs.cm−2. Duty cycle was locked to 100%. A total of 64 DIA_PASEF windows were used (25 Th isolation windows) with a cycle time of 1.8 seconds.

### Proteomics data analysis

For comparative analysis, we employed Spectronaut version 14.1 (Biognosys, Cambridge, MA) and utilized the default search unless otherwise indicated. All data were searched against the reviewed mouse proteome (UniProt UP000000589), using trypsin/LysC as the digestion enzymes. Cysteine carbamidomethylation was set as a fixed modification, while methionine oxidation and acetylation at the N-terminus were selected as variable modifications. A maximum of two missed cleavages and up to three variable modifications was allowed. The FDR cutoff was established at 1%, while the precursor peptide and q-value cutoffs were set at 0.2 and 0.01, respectively. In addition, protein q-value experiment and run-wide cutoffs were determined to be 0.01 and 0.05, respectively. Protein quantification was performed by filtering precursors using a q-value approach, using imputation with the background signal. The prototypicity filter was set to only protein group-specific peptides. Protein quantification was performed using MaxLFQ based on the area of MS2, with cross-run normalization enabled. Differential abundance testing was performed using unpaired t-tests and post-analysis based on both MS level on Spectronaut. In our analysis, we used a significance threshold of q value < 0.05, and an absolute average log2 (fold change) > 0.58 to identify significant candidates. Proteomics data was deposited in the Pride ProteomeXchange database (https://www.ebi.ac.uk/pride/), under the identifier PXD042593.

### Mitochondria isolation

Quadriceps muscle tissues were collected, flashed frozen in liquid nitrogen, and ground to powder in liquid nitrogen using a mortar and pestle. The powder was dissolved in an isolation Buffer (IB) containing: 0.22 M Mannitol, 0.07 M sucrose, 0.002 M Tris-HCL, 0.001 M EDTA, 0.4% BSA, and 20 mM Hepes at pH 7,5 and homogenized using a glass pestle in a 1.7 mL Microtubes (Axygen; Glendale, AZ,) with 10 strokes. The sample was then centrifuged 10 min at 800 g and the pellet was discarded. The supernatant was then centrifuged for 10 min at 14 000 g. The supernatant was discarded and the pellet containing the mitochondrial fraction was resuspended in IB.

### Co-immunoprecipitation

A day before the assay, 25 µl protein A magnetic beads S1425S (New England Biolabs; Ipswich, MA) were washed 3 times in PBS and incubated overnight with 2-3 µg of antibody directed against CISD3^4^ at 4°C with gentle mix. On the day of the assay, freshly isolated purified mitochondria were washed and pelleted for 10 min at 14 000 g. The mitochondria were resuspended in a lysis buffer containing 1% digitonin, 0.1% Noniped40, 0.15 M NaCl, 0.001 M EDTA, 0.05 M Hepes pH 7.5 and 0.00125 M dithiobis(succinimidyl propionate) (DSP) for 1 hr at 4°C with gentle mix. The mitochondria were then centrifuged at 14 000 g, the supernatant containing the protein extract collected, and protein content determined with The Pierce 660 nm Protein Assay (catalog number 1861426). 4mg/ml protein were then incubated with the coated beads with a gentle mix for four 4 hours at 4°C. The beads were there precipitated and washed five times with PBS and analyzed by MS and protein blots.

### Protein expression and purification

CISD3 (residues 36 to 127) and NDUFV2 (residues 32 to 249) cDNAs were inserted in the expression vector pet-28a+ (Novagen, Waltham, MA). Both proteins were expressed in Escherichia coli BL21-RIL grown in lysogeny broth supplemented with 30 μg/mL kanamycin and 34 μg/mL chloramphenicol. At an O.D. 600 nm of 0.6, the cells were supplemented with 0.75 mM FeCl3 (only for NDUFV2 protein), and protein expression was activated using 0.25 mM of isopropyl ß-D-1-thiogalactopyranoside. Cell growth proceeded for an additional 12 hr at 37 °C. NDUFV2 was purified from lysed cells using Ni-agarose and size exclusion chromatography as described^22,68^. CISD3 was purified by two consecutive rounds of ion exchange chromatography followed by size exclusion chromatography as described^4^. Apo-NDUFV2 was obtained by incubating the purified protein with EDTA and ferricyanide in the following molar ratio, 1:50:20; until the protein became colorless, and the protein was then dialyzed against 20 mM Tris pH 8.0 and 100 mM NaCl^75^.

### Protein-protein interaction

100 µM Holo-CISD3 and 100 µM Holo-NDUFV2, prepared as described above, were incubated in 20 mM Tris pH 8.0 and 100 mM NaCl for 10 min and the mixture was then denatured with or without β-mercaptoethanol for 6 hr. The samples were then separated using a 15% SDS-acrylamide gel. The band corresponding to the molecular weight of the complex CISD3-NDUFV2 was excised and analyzed by mass-spectrometry.

### [2Fe-2S] cluster transfer assay

Apo-NDUFV2 (250 µm) was pre-reduced in the presence of 5 mM Na-dithionite and 5 mM Na2-EDTA, pH 8.0, for 60 min. Apo-NDUFV2 was then incubated with holo-CISD3 (250 µm) in 50 mM Tris pH 8.0, 100 mM NaCl, 5 mM DTT, and 5 mM EDTA and [2Fe–2S] cluster transfer was analyzed by absorption spectroscopy from 380 nm to 700 nm at 37 °C as described in^4^. Data was acquired using a BioTek Synergy H1 Multimode Reader (Agilent, Santa Clara, CA).

### Protein-protein interaction prediction

A model for CISD3-NDUFV2 interaction was obtained after submission of the structure of CISD3 (PDB: 6AVJ) and NDUFV2 (extract from the structure of complex I PDB: 6G2J, chain E) to the web server HDOCK SERVER (hdock.phys.hust.edu.cn). The model had a docking score of −238.03 (knowledge-based iterative scoring function ITScorePP or ITScorePR; the protein-protein/RNA/DNA complexes in the PDB normally have a docking score of around −200) with a confidence score of 0.8533 (Confidence score = 1.0/[1.0+e0.02*(DockingScore+150)])^76,77^.

## Acknowledgments

This work was supported by the National Institute of Health grant GM111364 (to R.M.), the National Science Foundation (NSF)-Binational Science Foundation (BSF) Grant NSF-MCB 1613462 (to R.M.) and BSF Grant 2015831 (to R.N.), and The University of Missouri. We thank the Animal Modeling, Proteomics, Histopathology, and Electron Microscopy Research Core Facilities at the University of Missouri, Columbia.

## Author contribution

L.R, H-B.M., O.K., T.T.N., and D.G, performed the experiments and/or analyzed the data. R.M., H-B.M., C.M.A., and R.N. wrote the manuscript.

## Supplementary Materials

**Table S1:** Proteins with increased abundance in muscle tissue from *Cisd3^-/-^* mice.

**Table S2:** Proteins with decreased abundance in muscle tissue from *Cisd3^-/-^* mice.

**Fig. S1:** Weight of control and *Cisd3^-/-^*mice. Body weight of male (**a**) and female (**b**) control and *Cisd3^-/-^*mice at 13, 20 and 44 weeks is shown. Results are shown for male and female separately and presented as mean ± SD of 6 different animals (3 different males and 3 different females) from each group (WT and *Cisd3^-/-^* mice). Two-way ANOVA followed by a Tukey test was used to calculate statistical significance. White box and white square, WT male mice; Gray box and black square, *Cisd3^-/-^* male mice; white box and white circle, WT female mice; Gray box and black circle, *Cisd3^-/-^*female mice. Abbreviations: CISD, CDGSH Iron Sulfur Domain; WT, wild type.

**Fig. S2:** Centralized nuclei in muscle tissue from male of *Cisd3^-/-^* mice. **a** Representative image of cross sections of WT and *Cisd3^-/-^* male and female mice quadricep muscle stained with H&E. **b** Bar graph showing quantification of centralized nuclei in muscle tissue from male and female *Cisd3^-/-^*mice. Results are shown for male and female separately and presented as mean ± SD of 6 different animals (3 different males and 3 different females) from each group (WT and *Cisd3^-/-^* mice). Two-way ANOVA followed by a Tukey test was used to calculate statistical significance. White box and white square, WT male mice; Gray box and black square, *Cisd3^-/-^* male mice; white box and white circle, WT female mice; Gray box and black circle, *Cisd3^-/-^* female mice. Abbreviations: CISD, CDGSH Iron Sulfur Domain; WT, wild type.

**Fig. S3:** Abundance of different proteins involved in the DYSTROPHIN complex in *Cisd3^-/-^* mice compared to control wild type mice. Results are shown for male mice and presented as fold change compared to control. Two-way ANOVA followed by a Tukey test was used to calculate statistical significance. Abbreviations: CISD, CDGSH Iron Sulfur Domain; DMD, Duchenne muscular dystrophy; WT, wild type.

**Fig. S4:** Schematic diagrams of the wheel (**a**) and Rod (**b**) apparatuses used to measure grip strength of control and *Cisd3^-/-^* mice. Abbreviations: cm, centimeter.

