## Supplementary material for "CISD3 is required for Complex I function, mitochondrial integrity, and skeletal muscle maintenance": Suppl Figures

**Supplementary Table 1:** Protein data and statistical analysis of the proteomic comparison between male WT mice *vs* male *Cisd3*$^{-/-}$ mice.

**Supplementary Table 2:** Proteins significantly altered in their expression between male *WT* mice *vs* male *Cisd3*$^{-/-}$ mice.

**Supplementary Table 3:** List of all proteins belonging to each of the functional annotation groups presented in Fig. 5b.

**Supplementary Table 4:** Identity of proteins shown in the Venn diagram representing the comparison of proteins with significant altered expression from *Cisd3*$^{-/-}$ male *vs* other mice model.

**Supplementary Table 5:** Identity of proteins shown in the Venn diagram representing the comparison of mitochondrial proteins with significant altered expression from *Cisd3*$^{-/-}$ male *vs* other mice model.

**Supplementary Table 6:** Protein data analysis of the co-immunoprecipitation comparing male *WT* mice *vs* male *Cisd3*$^{-/-}$ mice presented in Fig. 7b.

**Supplementary Table 7:** Protein data analysis of the in-gel digestion presented in Fig. 8b.

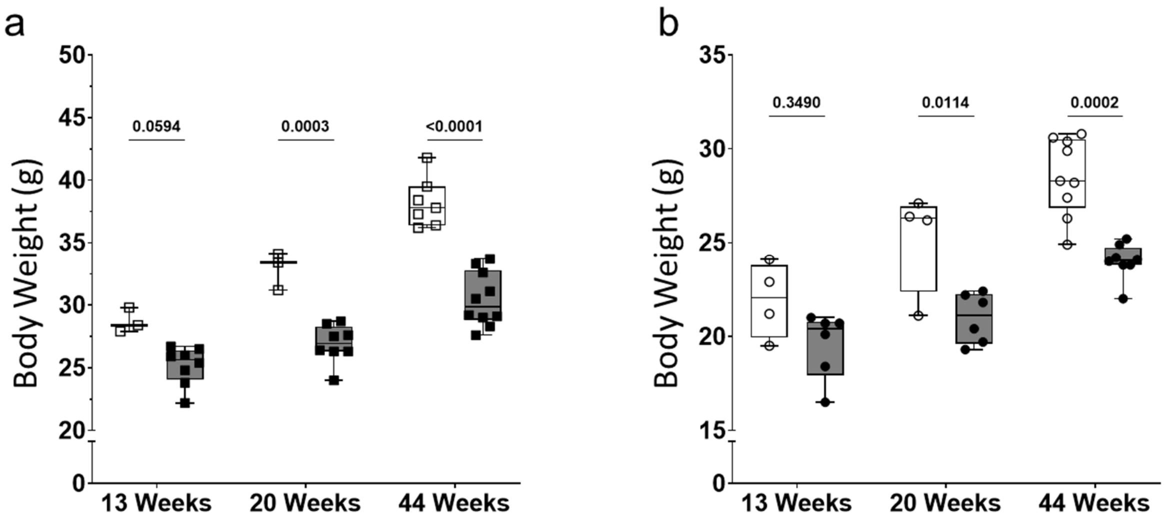


**Supplementary Figure 1:** Weight of control and *Cisd3*$^{-/-}$ mice. Body weight of male (**a**) and female (**b**) control and *Cisd3*$^{-/-}$ mice at 13, 20 and 44 weeks is shown. Results are shown for male and female separately and presented as box-and-whisker plots and include all data points of 6 different animals (3 different males and 3 different females) from each group (WT and *Cisd3*$^{-/-}$ mice). Two-way ANOVA followed by a Tukey test was used to calculate statistical significance. White box and white square, WT male mice; Gray box and black square, *Cisd3*$^{-/-}$ male mice; white box and white circle, WT female mice; Gray box and black circle, *Cisd3*$^{-/-}$ female mice. Abbreviations: CISD, CDGSH Iron Sulfur Domain; WT, wild type.

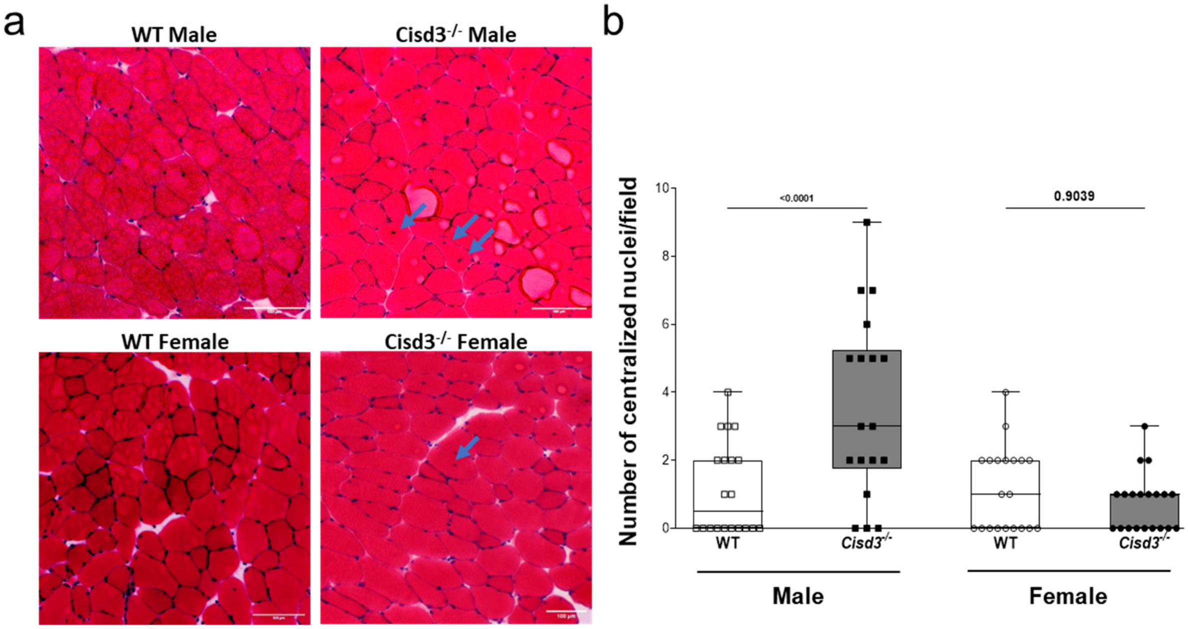


**Supplementary Figure 2:** Centralized nuclei in muscle tissue from male of *Cisd3*$^{-/-}$ mice. a Representative image of cross sections of WT and *Cisd3*$^{-/-}$ male and female mice quadricep muscle stained with H&E. b box-and-whisker plots showing quantification of centralized nuclei in muscle tissue from male and female *Cisd3*$^{-/-}$ mice. Results are shown for male and female separately and presented as box-and-whisker plots and include all data points of 6 different animals (3 different males and 3 different females) from each group (WT and *Cisd3*$^{-/-}$ mice). Two-way ANOVA followed by a Tukey test was used to calculate statistical significance. White box and white square, WT male mice; Gray box and black square, *Cisd3*$^{-/-}$ male mice; white box and white circle, WT female mice; Gray box and black circle, *Cisd3*$^{-/-}$ female mice. Abbreviations: CISD, CDGSH Iron Sulfur Domain; WT, wild type.

a

| gene name | DMD | *Cisd3*$^{-/-}$ | Uniprot ID | P-value | |
|---|---|---|---|---|---|
| Nos1 / Nitric Oxide Synthase 1 | 0.417385057 | 0.464829 | P82345 | 0.042989 | I |
| Dag1 / β-dystroglycan | 0.438828795 | 0.631179 | P82346 | 0.031354 | II |
| Sgcd / Delta-sarcoglycan | 0.357440319 | 0.57236 | P82347 | 0.043974 | III |

b

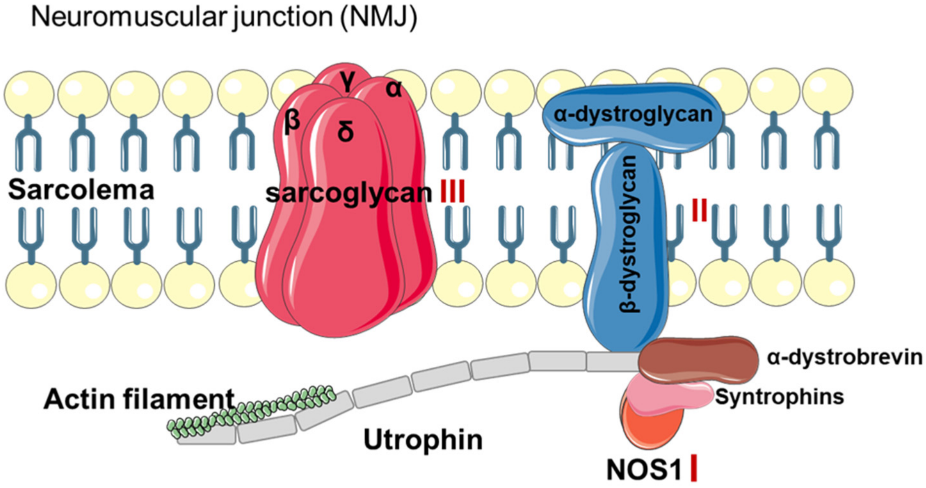


**Supplementary Figure 3:** Abundance of different proteins involved in the DYSTROPHIN complex in *Cisd3*$^{-/-}$ mice compared to control wild type mice. Results are shown for male mice and presented as fold change compared to control. Two-way ANOVA followed by a Tukey test was used to calculate statistical significance. Abbreviations: CISD, CDGSH Iron Sulfur Domain; DMD, Duchenne muscular dystrophy; WT, wild type.

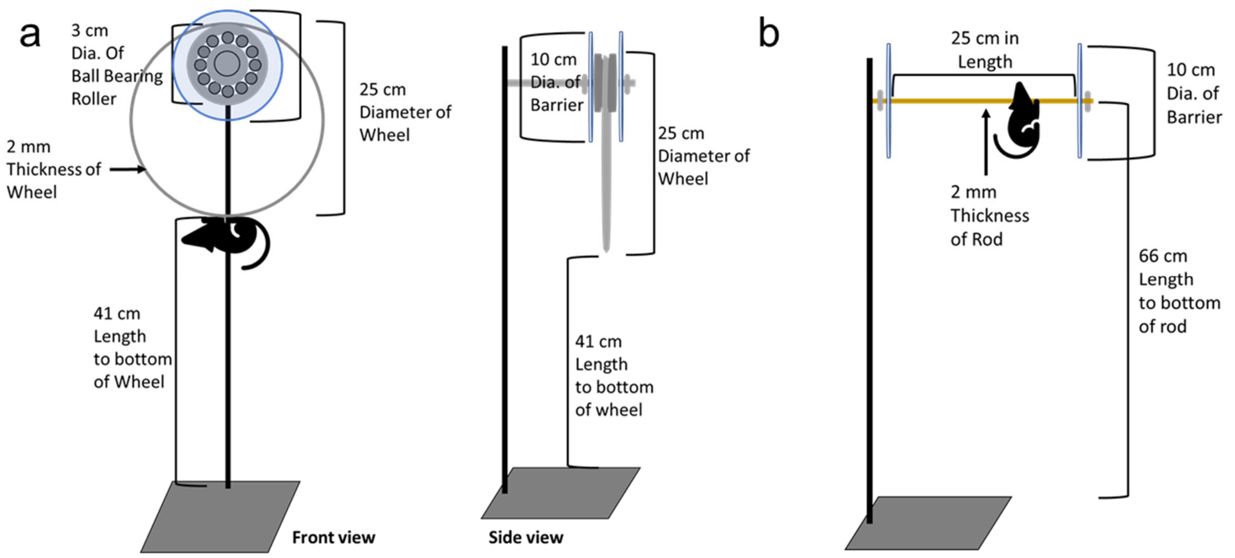


**Supplementary Figure 4:** Schematic diagrams of the wheel (a) and Rod (b) apparatuses used to measure grip strength of control and *Cisd3* mice. Abbreviations: cm, centimeter.
